# MetaboFM: A Foundation Model for Spatial Metabolomics

**DOI:** 10.1101/2025.10.23.684227

**Authors:** Efe Ozturk, Felix G. Rivera Moctezuma, Ahmet F. Coskun

**Author notes:** Corresponding author: Ahmet F. Coskun, Ph.D.

## Abstract

Mass spectrometry imaging (MSI) provides molecularly resolved maps of metabolites and lipids across tissues, yet the lack of large-scale, unified representation learning frameworks limits its potential for generalization and downstream analysis. Here, we introduce MetaboFM, a foundation model for spatial metabolomics that consolidates thousands of public MSI datasets into standardized spatial–spectral tensors and extracts transferable embeddings using pretrained Vision Transformers. We curated and standardized around 4000 publicly available MSI datasets from the METASPACE repository, spanning multiple organisms, tissue types, ionization modes, and instruments. Across six metadata prediction tasks—encompassing organism, ionization polarity, tissue type, condition, analyzer type, and ionization source embeddings from pretrained MetaboFM encoders achieved a mean macro–F1 of 0.74 and accuracy of 0.80 with linear probes, demonstrating substantially higher discriminative power than classical principal component analysis (PCA) or randomly initialized baselines by over 20 percentage points. To interpret the learned representations, we mapped embedding directions back to the *m/z* domain, revealing distinct spectral regions that drive class separation across tissues, conditions, and ionization sources. A multimodal visual question answering (VQA) extension further links MSI embeddings with natural-language queries through a cross-attention fusion module, attaining an average macro–F1 of 0.61 ± 0.05 across tasks. Finally, an interactive Gradio interface enables users to visualize MSI patches and query sample metadata in free-form language. Together, MetaboFM establishes a scalable foundation model paradigm for MSI, unifying representation learning, spectral interpretability, and multimodal interaction within a single framework for spatial metabolomics.

## 1 Introduction

MSI enables direct, label-free mapping of the spatial distribution of metabolites, lipids, and proteins across biological tissues[1, 2, 3]. Unlike conventional microscopy-based approaches that rely on fluorescent or chromogenic labeling, MSI provides molecularly resolved ion distributions that can span thousands of channels, offering an unbiased view of tissue chemistry and metabolism. However, MSI data are inherently high-dimensional, often noisy, and sparsely annotated. Differences in sample preparation, ionization modality, and instrumentation further introduce substantial variability across datasets. As a result, MSI studies are typically analyzed in isolation, and the lack of unified representation learning frameworks limits generalization and reuse of data at scale.

In recent years, foundation models have revolutionized representation learning in computer vision, natural language processing, and single-cell omics by pretraining large neural networks on unlabeled data to obtain general-purpose embeddings[4, 5, 6, 7]. Approaches such as CLIP and DINOv2 demonstrate that self-supervised pretraining can yield transferable visual representations that generalize across tasks and domains[5, 6]. Within spatial omics, analogous paradigms are beginning to emerge—such as Kronos and scFoundation—which apply self-supervised learning to cellular or molecular images[7, 8]. Yet, a comparable foundation model dedicated to MSI has not been established. Existing pipelines remain highly specialized, reliant on handcrafted preprocessing steps, or restricted to supervised learning on limited labeled data[9, 10].

Here, we introduce MetaboFM, a foundation model framework tailored for MSI. MetaboFM unifies standardized data curation and multimodal representation within a cohesive architecture. The framework consolidates MSI tiles from a diverse public repository METASPACE [11] into consistent spatial–spectral tensors and employs pretrained Vision Transformer (ViT) backbones—such as DINOv2, DeiT, and MAE—to derive robust feature embeddings (Figure 1a)[6, 12, 13]. These embeddings capture spatial organization and molecular composition across modalities, enabling broad generalization across tissue types, organisms, and acquisition conditions.

**Figure 1:**
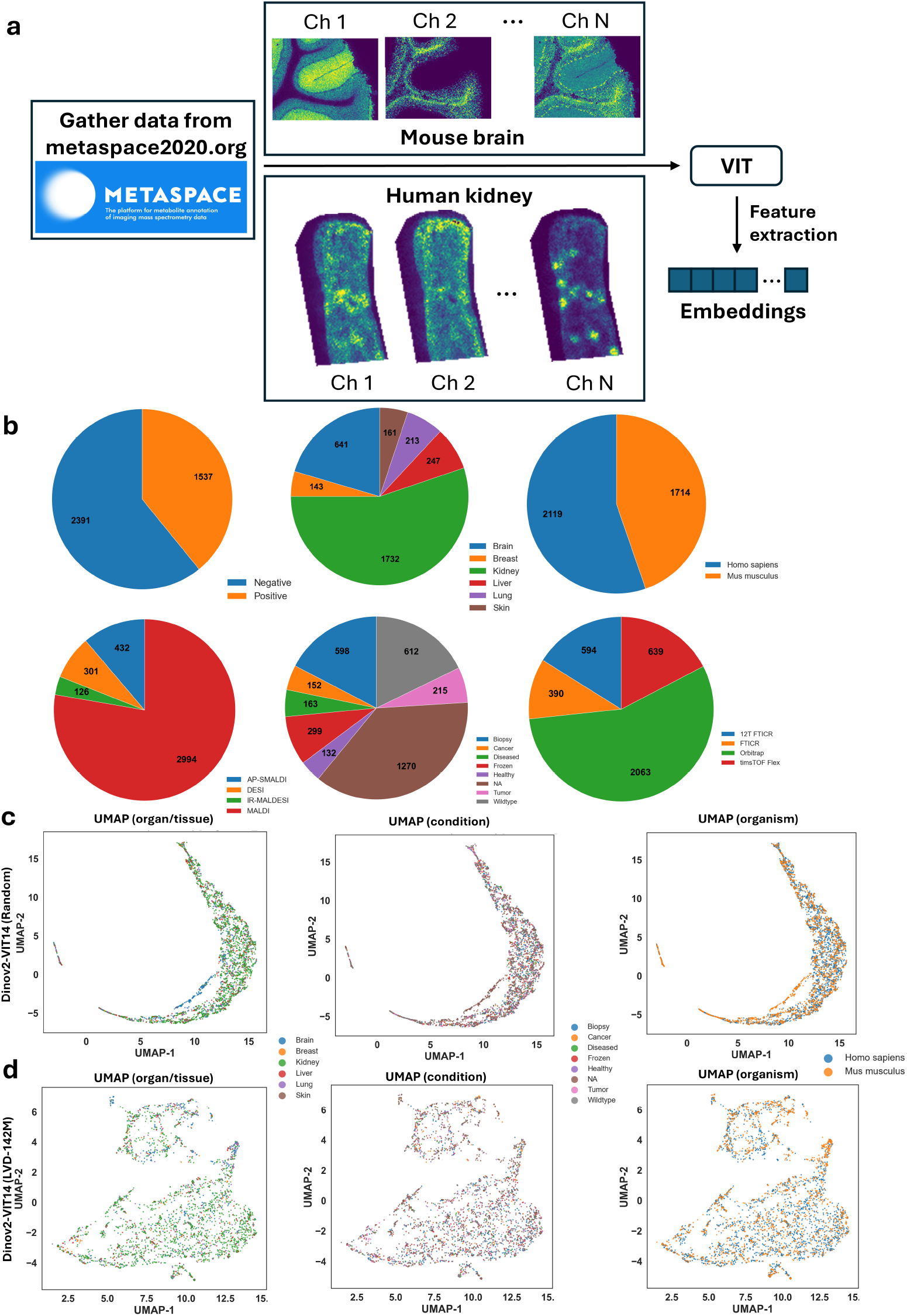
Overview of data collection, feature extraction, and representation analysis in MetaboFM. **(a)** Schematic of the data gathering and feature extraction workflow. MSI datasets are collected from the METASPACE repository (https://metaspace2020.org), processed into standardized multi-channel tiles, and passed through a pretrained ViT encoder to obtain spatial–chemical embeddings. **(b)** Class distribution across metadata prediction tasks, illustrating the diversity and imbalance of samples among categories for *organism, ionization polarity, organ/tissue, condition, analyzer type*, and *ionization source*. **(c)** Two-dimensional UMAP projections of embeddings produced by a randomly initialized DINOv2–ViT–B/14 model for three representative tasks (*organ/tissue, condition*, and *organism*). **(d)** Corresponding UMAP projections generated using the pretrained DINOv2–ViT–B/14 (LVD-142M) encoder.

To extend MetaboFM beyond representation learning, we integrate a VQA component that links the pretrained MSI encoder with a lightweight text encoder via cross-attention. This module enables natural-language queries—such as “What organism is this sample?” or “What is the ionization polarity?”—and returns categorical predictions grounded in the learned MSI representations[5, 14]. Finally, we deploy the framework through a Gradio-based graphical interface that allows researchers to upload MSI patches, visualize spatial distributions, and interactively query metadata, making the system accessible to the broader spatial omics community[15].

To benchmark the generality of MetaboFM across diverse biological and experimental contexts, we evaluated six categorical metadata prediction tasks encompassing *organism, ionization polarity, organ/tissue, condition, analyzer type*, and *ionization source* [16]. These categories span examples such as human versus mouse specimens, positive versus negative ionization modes, liver versus brain tissues, healthy versus diseased conditions, and distinct mass spectrometer analyzers. The overall distribution of samples across these classes highlights both the scale and imbalance of the collected corpus (Figure 1b).

Together, MetaboFM establishes a general-purpose foundation model paradigm for MSI, bridging the gap between raw ion distributions and interpretable biological information. The framework standardizes data processing and representation extraction, providing a unified pipeline for large-scale MSI analysis and multimodal integration. In this study, we evaluate MetaboFM across multiple pretrained Vision Transformer architectures to assess the transferability and robustness of visual representations for MSI data. By unifying standardized data curation, pretrained feature extraction, and multimodal querying, MetaboFM offers a scalable foundation for future integration of metabolomic, histological, and proteomic imaging modalities[8, 17].

## 2 Results

### 2.1 Representation Learning and Visualization

To evaluate the representational structure of MSI embeddings, we first examined UMAP projections of DINOv2–ViT–B/14 encoders with and without pretraining. Randomly initialized embeddings showed diffuse and overlapping distributions across metadata categories, whereas pretrained DINOv2–ViT–B/14 (LVD-142M) embeddings exhibited compact and well-separated clusters corresponding to biological attributes such as *organ/tissue, condition*, and *organism* (Figure 1c–d). These results demonstrate that large-scale visual pretraining substantially improves the organization and separability of MSI latent spaces.

### 2.2 Linear Probe Evaluation

We next assessed the discriminative power of embeddings using linear probes trained on all available training data (Figure 2a). The pretrained DINOv2–ViT–B/14 (LVD-142M) achieved the highest overall macro–F1 across tasks (0.7410; accuracy 0.7963), followed by DeiT-B/16 (0.7370; 0.7912) and MAE–ViT–B/16 (0.7170; 0.7766), while randomly initialized variants averaged around 0.5513 (0.6069 accuracy). The PCA baseline performed lowest (0.4742; 0.5264). Across individual tasks, DINOv2 achieved the top performance for *condition* (0.6369; 0.6984), *analyzer type* (0.7990; 0.8415), and *polarity* (0.8206; 0.8292), whereas DeiT-B/16 excelled in *organ/tissue* (0.6919; 0.7896) and *ionization source* (0.7826; 0.8853), and MAE–ViT–B/16 was best for *organism* (0.8184; 0.8184). These results demonstrate that self-supervised pretraining, particularly with large-scale DINOv2 embeddings, yields transferable representations that generalize across diverse acquisition modalities and metadata categories.

**Figure 2:**
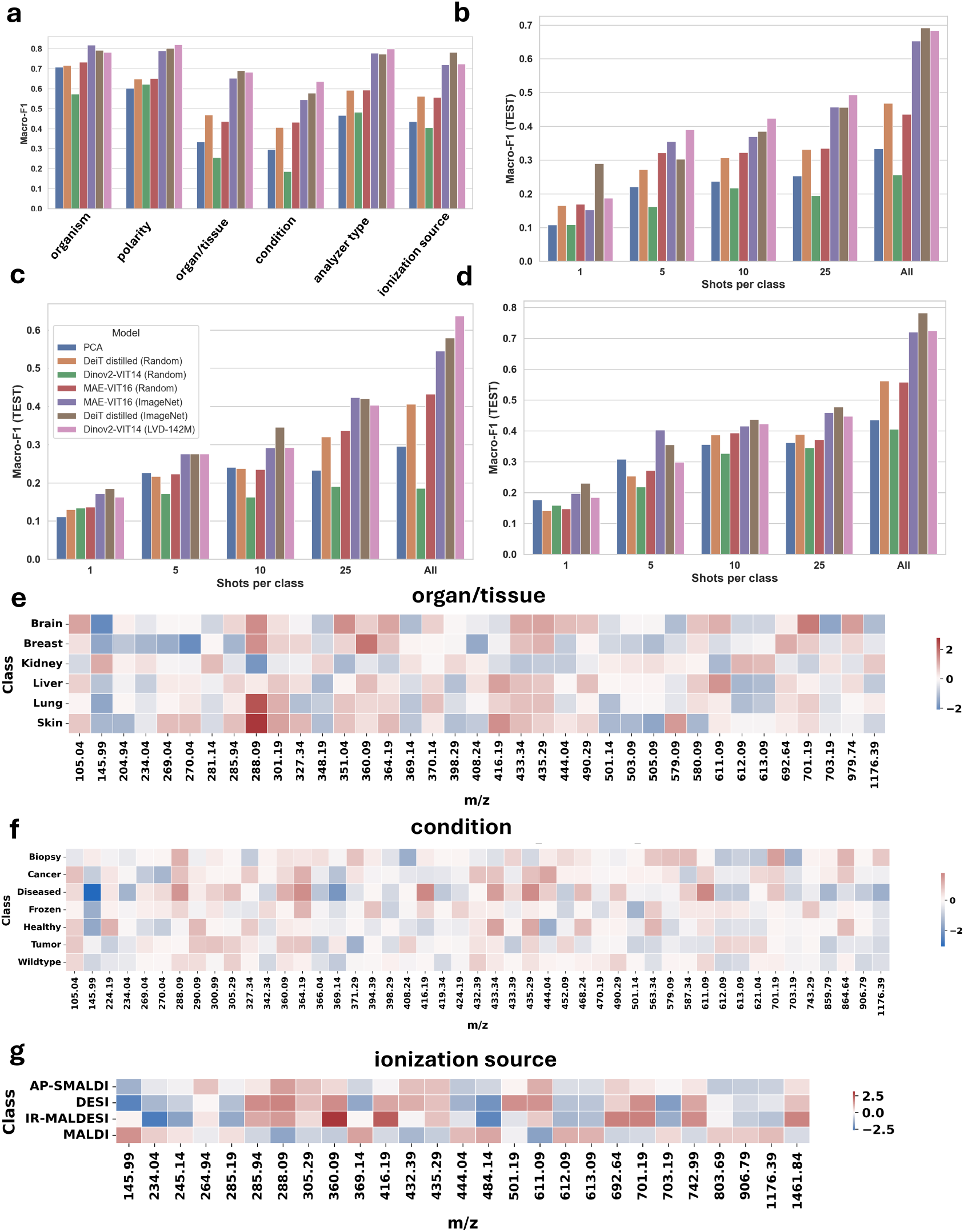
Downstream performance and spectral attribution of the MetaboFM framework. **(a)** Macro–F1 across metadata tasks using linear probes trained on all available data. **(b–d)** Few-shot macro–F1 performance for *organ/tissue, condition*, and *ionization source* tasks. **(e–g)** m/z importance maps for the same three tasks, computed by projecting class-specific embedding directions onto the binned spectral domain (**s**^(*c*)^ = **Bw**^(*c*)^). Red and blue indicate positive and negative contributions of individual *m/z* bins toward class discrimination, highlighting molecular regions most associated with each category.

We further evaluated few-shot transfer by training linear probes with {1, 5, 10, 25} labeled samples per class across representative metadata tasks (Figure 2b–d). For the *condition* task, DINOv2 achieved macro–F1 of 0.16, 0.28, 0.29, and 0.41 for 1-, 5-, 10-, and 25-shot settings, respectively, reaching 0.64 when trained on all samples. For *organ/tissue*, DINOv2 reached 0.19, 0.39, 0.42, and 0.50 across the same shot counts, attaining 0.68 with all data. For *ionization source*, performance rose from 0.19 to 0.73 across the few-shot spectrum. In all cases, randomly initialized encoders and PCA remained below 0.40 at low shot counts.

### 2.3 Spectral Attribution and m/z Importance

To interpret which molecular features drive class separation, we projected embedding-based class directions back to the binned *m/z* axis using the surrogate mapping described in Section 3.4. The resulting attribution heatmaps revealed distinct spectral regions contributing to each class in the *organ/tissue, condition*, and *ionization source* tasks (Figure 2e–g).

For *organism/tissue*, discriminative *m/z* intervals included approximately 285.94–327.34, and 416.19–490.29, with elevated contributions for skin and lung tissues around *m/z* 288.09 (Figure 2e). For *condition*, strongly weighted bins appeared near *m/z* 290.09–305.29, highlighting molecular differences between *healthy*, and *tumor* states (Figure 2f). For *ionization source*, the dominant ranges were found around *m/z* 285.94–435.29 (Figure 2g).

### 2.4 Visual Question Answering Framework

We extended MetaboFM with a VQA module that integrates MSI embeddings from the Vision Transformer and natural-language inputs from a text encoder through a cross-attention fusion module (Figure 3a). Across six metadata tasks, the model achieved robust five-fold cross-validation performance with an average macro–F1 of 0.61±0.05 and accuracy of 0.74±0.03 (Figure 3b). Performance varied by task, reaching macro–F1 values of 0.74 for *organism*, 0.71 for *polarity*, 0.68 for *analyzer type*, 0.57 for *ionization source*, 0.55 for *organ/tissue*, and 0.41 for *condition*. The corresponding accuracies followed a similar trend, peaking at 0.88 for *ionization source* and remaining above 0.63 for all major categories. These results confirm that the multimodal fusion architecture effectively aligns image-derived spatial–chemical features with linguistic queries, enabling accurate metadata reasoning across diverse MSI datasets.

**Figure 3:**
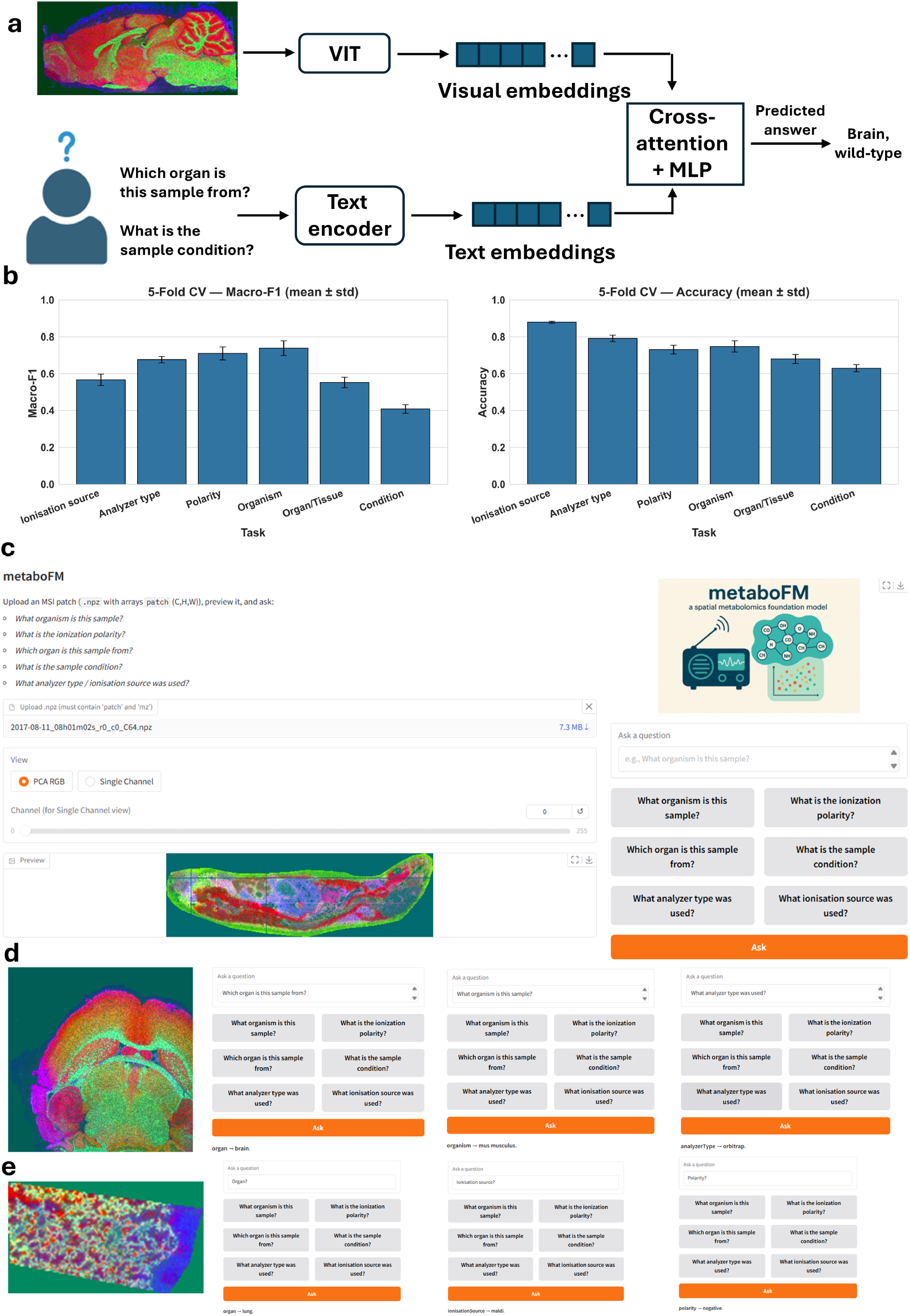
Visual question answering and interactive interface of MetaboFM. **(a)** Schematic of the VQA framework linking MSI embeddings from a Vision Transformer with a text encoder through a cross-attention fusion module. **(b)** Five-fold cross-validation results showing macro–F1 and accuracy across metadata tasks. **(c)** Overview of the Gradio-based graphical user interface for interactive querying and visualization of MSI patches. **(d)** Example MSI sample loaded into the interface with predictions for predefined question templates. **(e)** Demonstration on another sample using a custom free-form question, illustrating flexible natural-language interaction with the trained VQA model.

### 2.5 Interactive Interface and Example Outputs

We deployed the trained VQA model in a Gradio-based graphical interface enabling direct interaction with MSI data. Users can upload MSI patches, visualize them in PCA-RGB or single-channel modes, and query the model using predefined or free-form questions (Figure 3c). Example interactions show accurate predictions for structured templates such as “What organism is this sample?” (Figure 3d) as well as flexible responses to custom questions like “Polarity?” (Figure 3e), demonstrating the accessibility and interpretability of MetaboFM for interactive spatial metabolomics analysis.

## 3 Methods

### 3.1 Data Collection and Preprocessing

MSI datasets were aggregated from publicly available repositories, primarily METASPACE, focusing on human and mouse tissue studies[11, 16]. Each dataset contains molecular annotations linked to ion images representing the spatial distributions of detected metabolites or lipids. To ensure consistency and coverage across studies, we standardized the curation pipeline to process every dataset under identical quality control and sampling criteria before model training.

The first stage involved filtering molecular annotations by a strict false discovery rate (FDR) threshold of 10%, ensuring high-confidence assignments while maintaining sufficient dataset diversity[18]. When available, annotations were ranked by the Metabolite Signal Match (MSM) score, reflecting annotation confidence and isotopic pattern quality. This ranking helped prioritize informative channels, and a fixed cap was placed on the number of annotations retained per dataset to avoid overrepresentation by large studies. For each annotation, we extracted up to three isotope images corresponding to the monoisotopic peak and its first two isotopologues (M, M+1, and M+2), balancing information retention with manageable storage requirements.

Each ion image was normalized through percentile-based intensity scaling. Specifically, the 1st and 99th percentile values of the image intensity distribution were used to remove extreme outliers and harmonize contrast across datasets. Intensities were rescaled to the [0, 1] range and stored as 16-bit unsigned integers, substantially reducing storage and improving loading performance while preserving spatial intensity patterns. To ensure consistent spatial context, all images were tiled into fixed non-overlapping squares of 256 × 256 pixels. Tiles with fewer than 3% nonzero pixels were automatically excluded to remove near-blank regions or off-tissue areas. Basic per-tile statistics such as mean, standard deviation, and nonzero fraction were recorded for downstream quality control and filtering.

Because instrument and acquisition metadata in public repositories are often inconsistent, a harmonization step was introduced. This stage parsed the original metadata files and standardized key experimental attributes—organism, tissue or organ part, condition, analyzer type, ionization source, and ionization polarity—using a case-insensitive, variant-aware resolver. This approach consolidated heterogeneous naming conventions (e.g., “MS Analysis” vs. “MS Analysis”) into a unified schema, facilitating cross-dataset compatibility and later stratified evaluations[16] (see Table 1 for the distribution of samples across metadata tasks).

**Table 1:** Dataset summary showing the number of samples per class across six metadata prediction tasks. Counts are shown in parentheses.

| Condition | Organism Part | Analyzer Type | Ionisation Source | Organism | Polarity |
| --- | --- | --- | --- | --- | --- |
| NA (1270) | Kidney (1732) | Orbitrap (2063) | MALDI (2994) | Homo sapiens (2119) | Negative (2391) |
| Wildtype (612) | Brain (641) | timsTOF Flex (639) | AP-SMALDI (432) | Mus musculus (1714) | Positive (1537) |
| Biopsy (598) | Liver (247) | 12T FTICR (594) | DESI (301) | Mixed (72) |  |
| Frozen (299) | Lung (213) | FTICR (390) | IR-MALDESI (126) |  |  |
| Tumor (215) | Skin (161) | TOF reflector (73) |  |  |  |
| Diseased (163) | Breast (143) |  |  |  |  |
| Cancer (152) | Heart (65) |  |  |  |  |
| Healthy (132) | Fibroblasts (56) |  |  |  |  |
|  | Ovary (55) |  |  |  |  |
|  | Whole organism (52) |  |  |  |  |

The processed ion images were then grouped into multi-channel tiles suitable for feature extraction. For each dataset, all images corresponding to the same spatial coordinates were collected and stacked into a tensor representing a fixed field of view with multiple molecular channels. Up to sixty-four channels were included per tile, prioritized by annotation rank to maximize information density, while groups with fewer than eight channels were excluded. Each channel stack was stored alongside its associated *m/z* values, ensuring consistent linkage between spatial and molecular information.

Finally, the dataset was partitioned into training and testing subsets using a deterministic hash function applied to dataset identifiers. This procedure ensured that all tiles originating from the same dataset were assigned to a single split, preventing information leakage and maintaining reproducibility. Together, these steps produced a standardized, high-quality corpus of MSI tiles spanning multiple organisms, instruments, and acquisition conditions—serving as the foundation for large-scale representation extraction and downstream evaluation within the MetaboFM framework.

### 3.2 Foundation Model Architecture and Feature Extraction

MSI data differ fundamentally from conventional natural images: each pixel encodes a molecular spectrum across numerous *m/z* channels rather than color intensities. To derive transferable representations from these spatial–chemical patterns, MetaboFM leverages pretrained ViT architectures originally trained on large-scale natural image datasets[4]. These encoders act as general-purpose feature extractors that capture spatial organization and texture features relevant to ion-distribution patterns in MSI.

Let **X** ∈ ℝ^*C*×*H*×*W*^ denote an MSI tile containing *C* ion images (molecular channels). Each channel **X**_*c*_ ∈ ℝ^*H*×*W*^ is normalized and replicated across three channels to form a pseudo-RGB image

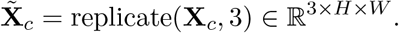

The replicated image is resized to match the input resolution of a pretrained ViT backbone *f*_*θ*_. The model processes 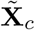 and outputs a high-level representation corresponding to the *class token (CLS)* embedding, a learned global feature that summarizes information from all image patches:

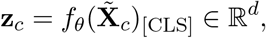

where **z**_*c*_ encodes the global spatial features of the ion channel image.

For each MSI tile, the per-channel embeddings are aggregated to form a single representation vector by taking their mean and applying *l*_2_ normalization:

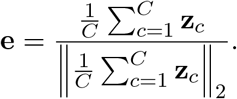

This aggregated embedding **e** ∈ ℝ^*d*^ provides a compact and modality-agnostic descriptor of the molecular and structural context of each MSI tile.

MetaboFM supports multiple pretrained encoders, including ImageNet-supervised DeiT-B/16, self-distilled DINOv2 ViT-S/14 and ViT-B/14, and the MAE ViT-B/16 encoder. These models remain fixed during feature extraction, allowing systematic comparison of how large-scale visual pretraining transfers to molecular imaging data.

### 3.3 Downstream Evaluation

#### 3.3.1 Tasks and Metrics

We evaluate six categorical metadata prediction tasks: *organism, ionization polarity, organ/tissue, analyzer type, ionization source*, and *condition*. The primary evaluation metric is macro–F1 on the held-out test split, which accounts for class imbalance across categories.

#### 3.3.2 Data Splits and Preprocessing

To prevent information leakage, training and test sets are defined at the *dataset* level—ensuring that samples from the same dataset do not appear in multiple splits. Each MSI patch is center-cropped or padded to a fixed spatial size and reduced to a consistent number of channels using a variance-biased selection with anchor-channel repetition when necessary. This preprocessing mirrors the configuration used during embedding extraction.

#### 3.3.3 Embeddings and Feature Extraction

For all experiments, embeddings are extracted directly from pretrained Vision Transformer backbones. Each MSI tile is normalized, converted into a pseudo-RGB format per ion channel, and processed by the pretrained encoder to obtain per-channel features that are averaged into a single *l*_2_-normalized embedding vector. No fine-tuning is performed. These embeddings serve as fixed inputs for all downstream evaluators. While embeddings are generated for all splits, the test set is reserved strictly for final reporting.

#### 3.3.4 Models Compared

We benchmark a diverse set of Vision Transformer architectures and baselines reflecting varying initialization and pretraining strategies. These include a pixel-level baseline based on PCA; randomly initialized models including the Data-efficient Image Transformer with 16-pixel patches(*DeiT-B/16*), the self-distilled DINO Vision Transformer with 14-pixel patches (*DINOv2–ViT–B/14*), and the Masked Autoencoder Vision Transformer with 16-pixel patches (*MAE–ViT–B/16*); ImageNet-supervised models comprising the *MAE–ViT–B/16* and *DeiT-B/16* variants; and the large-scale self-distilled foundation model trained on 142 million natural images (*DINOv2–ViT–B/14 (LVD-142M)*). All models share identical preprocessing, normalization, and embedding extraction pipelines to ensure fair and consistent comparison across architectures.

#### 3.3.5 Probing Protocols

To quantify downstream task performance, we train a multinomial logistic-regression probe with standardized input features. To evaluate label efficiency, we perform few-shot experiments using 1, 5, 10, 25 labeled samples per class drawn uniformly from the training split, alongside a full-data condition utilizing all available training examples.

### 3.4 m/z Importance Analysis

To interpret how individual molecular features contribute to downstream categorical predictions, we established an attribution analysis linking the spectral domain of the MSI data to the latent representations derived by the foundation model [19]. This procedure estimates how specific *m/z* values influence model embeddings and, in turn, task-level class separation, thereby providing a continuous map from spectral content to semantic output space.

Let each MSI tile be represented by its raw intensity spectrum **x**(*m*) defined over a set of centroided *m/z* values. To obtain a consistent representation across all tiles, the spectra were projected onto a common *m/z* grid by discretizing the full observed range into uniform bins of width Δ*m* = 0.05 Da. The resulting binned intensity vector for tile *i* is denoted **x**_*i*_ ∈ ℝ^*M*^, where *M* is the number of discrete *m/z* bins. These vectors collectively form the spectral matrix **X**_*mz*_ ∈ ℝ^*N* ×*M*^ for all *N* analyzed tiles. The corresponding embedding matrix **Z** ∈ ℝ^*N* ×*D*^ contains the *D*-dimensional latent representations previously extracted from the pretrained vision encoder.

We then sought a linear correspondence between spectral intensities and embedding coordinates by solving a regularized least-squares problem,

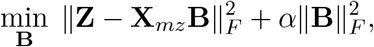

where *α* is a ridge regularization parameter. The matrix **B** ∈ ℝ^*M* ×*D*^ captures how variations at each *m/z* bin relate to directions in the embedding space. Each column of **B** thus represents the spectral signature associated with one latent dimension of the pretrained model.

To relate embeddings to semantic categories, a sparse linear model was fit independently for each downstream classification task defined in Section 3.3. For a given task *t* with class set *C*_*t*_, a weight vector **w**^(*c*)^ ∈ ℝ^*D*^ was obtained for every class *c* ∈ *C*_*t*_ such that **w**^(*c*)^ describes the direction in the embedding space that maximally distinguishes class *c* from others. These vectors quantify how class membership varies as a linear function of the pretrained representations.

By combining the two mappings, we obtain a continuous attribution of each *m/z* bin to every semantic class through

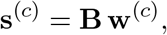

where **s**^(*c*)^ ∈ ℝ^*M*^ assigns a signed importance score to each *m/z* position. Positive values indicate bins whose spectral intensity contributes positively to the latent direction associated with class *c*, whereas negative values indicate inhibitory or opposing influence. The resulting importance matrix 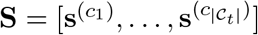 provides a direct bridge between molecular features and categorical semantics learned by the foundation model.

For visualization, the most informative *m/z* bins were identified by selecting those with the largest absolute scores across all classes. The corresponding importance values were normalized and displayed as a two-dimensional heatmap, with rows representing classes and columns representing *m/z* bins. This representation reveals which molecular mass regions are most strongly associated with specific biological categories within the latent space, offering a mechanistic interpretation of the model’s learned representations.

### 3.5 Visual Question Answering Framework

We extend the foundation model with a VQA module that maps natural-language questions to *categorical* metadata predictions (organism, ionization polarity, organ/tissue, analyzer type, ionization source, and condition)[5, 14]. The module conditions answers on the model’s spatial–chemical token representations while keeping the backbone frozen.

An MSI patch is encoded by the pretrained vision encoder (*DINOv2–ViT–B/14 (LVD-142M)*) into a set of visual tokens in ℝ^*d*^. A question is encoded by a text encoder (*Sentence-Transformers all-MiniLM-L6-v2*) [20] into a single *d*-dimensional embedding produced by mean pooling over the final Transformer layer outputs, followed by *l*_2_ normalization to match the vision embedding space. Language–vision fusion is performed by a single cross-attention block in which the question acts as the query and the visual tokens as keys/values:

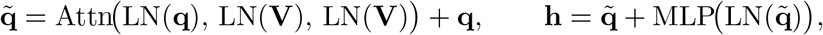

where Attn is multi-head attention, LN denotes layer normalization, and the MLP is a two-layer GELU block with a residual connection. The fused representation **h** ∈ ℝ^*d*^ is passed to task-specific linear classifiers. For each categorical task *t* with label set **C**_*t*_, logits are computed as **o**^(*t*)^ = **W**_*t*_**h** and converted to probabilities by softmax(**o**^(*t*)^).

Free-form questions are routed to the appropriate classifier via a lightweight intent matcher that recognizes domain terms (e.g., “organism”, “polarity”, “organ/tissue”, “analyzer type”, “ionization source”, “condition”). This preserves a natural interface while ensuring supervision and evaluation are applied to the correct prediction space.

Training pairs each MSI patch with one or more metadata-templated questions (e.g., “What organism is this sample?”). Let *t*_*b*_ be the routed task for question *b* and 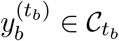 its groundtruth label. The objective sums cross-entropy over only the applicable categorical heads in the mini-batch:

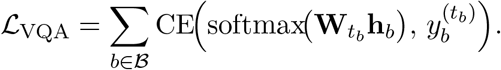

The MSI encoder remains frozen to preserve pretrained semantics; only the text encoder, crossattention block, and task heads are updated. We use AdamW, gradient-norm clipping, and track an exponential moving average of the training loss.

To standardize inputs, patches are center-cropped/padded to a fixed size and reduced to a small, fixed number of channels via a variance-biased selection with anchor-channel repetition when needed. This mirrors the foundation model’s view sampling and yields robustness to missing ion images and heterogeneous channel counts.

In deployment, the system accepts an MSI patch and a free-form question, performs intent routing, and returns the corresponding categorical answer with confidence (and optional top-*k*). Design choices—single cross-attention layer for low latency, a frozen backbone to avoid overfitting, per-task classifiers to prevent negative transfer, and lightweight intent detection—combine to provide a simple, fast, and well-grounded VQA interface for MSI metadata queries.

### 3.6 Graphical User Interface with Gradio

A lightweight Gradio web application provides an interactive interface for qualitative evaluation and user studies. The app allows users to upload MSI patches stored as.npz files, preview them in PCA-RGB or single-channel mode, and ask free-form questions such as “What organism is this sample?” or “What is the ionization polarity?”. Each question is automatically routed to the appropriate categorical head, and the predicted answer with confidence score is displayed alongside the MSI preview. The interface also includes quick-pick question buttons and adjustable viewing parameters, enabling accessible exploration of the VQA model’s predictions across metadata domains.

## 4 Discussion and Conclusion

MetaboFM establishes a unified foundation model for MSI, demonstrating that large-scale visual pretraining can produce transferable and interpretable representations for spatial metabolomics. The pretrained VIT encoders yielded structured embeddings that generalize across biological and experimental conditions, achieving superior discriminative power compared to PCA and randomly initialized baselines. The VQA module further linked these embeddings to natural-language queries, enabling interactive metadata reasoning through a graphical interface.

The spectral attribution analysis revealed that specific *m/z* intervals drive class separation across tissues, conditions, and ionization sources, providing interpretable molecular insights from embedding spaces. This connection between learned representations and chemical specificity highlights the potential of foundation models to uncover biologically meaningful patterns within complex MSI datasets.

Despite these advances, challenges remain. Heterogeneity and limited standardization across public MSI repositories may introduce dataset bias, and the current framework relies on frozen backbones without domain-specific pretraining. Future work will explore self-supervised and multimodal pretraining directly on MSI data, extending MetaboFM to histological and proteomic modalities for comprehensive spatial omics integration.

Building upon recent progress in guided super-resolution approaches that integrate high-resolution imaging mass cytometry with MSI to achieve single-cell–level spatial metabolomics [21], the next step is to unify these advances within foundation model frameworks. MetaboFM moves toward this direction by providing scalable, generalizable representations that can ultimately enable single-cell metabolomic analysis at the tissue scale.

In summary, MetaboFM presents a large-scale foundation model framework for MSI, combining representation learning, spectral interpretability, and multimodal interaction. It lays the groundwork for next-generation computational metabolomics by bridging molecular imaging, machine learning, and biological understanding.

## Supporting information

MetaboFM Supplemental Movie

## 5 Code availability

All code for data curation, feature extraction, representation learning, visual question answering, and spectral attribution is available in the MetaboFM repository at https://github.com/coskunlab/MetaboFM.

